# Aversive Learning Increases Release Probability of Olfactory Sensory Neurons

**DOI:** 10.1101/642751

**Authors:** Janardhan P. Bhattarai, Mary Schreck, Andrew H. Moberly, Wenqin Luo, Minghong Ma

## Abstract

Predicting danger from previously associated sensory stimuli is essential for survival. Contributions from altered peripheral sensory inputs are implicated in this process, but the underlying mechanisms remain elusive. Here we use the mammalian olfactory system to investigate such mechanisms. Primary olfactory sensory neurons (OSNs) project their axons directly to the olfactory bulb (OB) glomeruli where their synaptic release is subject to local and cortical influence and neuromodulation. Pairing optogenetic activation of a single glomerulus with foot shock in mice induces freezing to the light stimulation alone during fear retrieval. This is accompanied by an increase in OSN release probability and a reduction in GABA_B_ receptor expression in the conditioned glomerulus. Furthermore, freezing time is positively correlated with the release probability of OSNs in fear conditioned mice. These results suggest that aversive learning increases peripheral olfactory inputs at the first synapse, which may contribute to the behavioral outcome.

## Introduction

Associative learning, the process by which the brain assigns predictive values to sensory stimuli and adjusts behavioral responses, is essential for survival. This process is mediated by neuroplasticity along the sensory pathways as well as in associated cortical and limbic regions [1–3]. Although contributions from altered peripheral sensory inputs into the brain have been implicated [4–7], the underlying mechanisms remain poorly understood.

The mammalian olfactory system provides an experimentally accessible circuit to study learning-induced plasticity in relation to behavioral output. Odor detection and discrimination rely on a large family (>1000 in rodents) of G protein-coupled odorant receptors (ORs) expressed in olfactory sensory neurons (OSNs) in the nasal epithelium. Each OSN expresses a single OR type and all the OSNs with the same OR identity project their axons typically onto two glomeruli (one medial and one lateral) in each olfactory bulb (OB) [8, 9]. The axon terminals of OSNs make synaptic contacts with both glutamatergic projection neurons (mitral and tufted cells) and GABAergic interneurons in the OB. In the glomerular layer external tufted (ET) cells receive monosynaptic inputs from OSNs, while short-axon and periglomerular (PG) cells receive either direct or indirect OSN inputs [10, 11]. The PG cells provide presynaptic inhibition onto OSNs via GABA_B_ receptors at the axonal terminals [12–17]. As PG cells also receive top-down centrifugal inputs, transmitter release from OSNs is subject to both local and cortical influence and neuromodulation [18, 19]. This strategy of modulating sensory inputs into the brain via presynaptic GABA_B_ receptors and local inhibitory neurons is widely adapted by other sensory systems (e.g., visual and auditory) as well [20, 21].

The OSN axon terminals release glutamate onto postsynaptic cells and the release properties can be accessed by recording excitatory postsynaptic currents (EPSCs) in the postsynaptic cells. Olfactory nerve stimulation induces a form of short term plasticity called paired pulse depression; i.e. the EPSC evoked by the second pulse is smaller than the first one. The paired pulse ratio (PPR) of evoked EPSCs (second/first) is routinely used as indication of release probability: a higher (or lower) PPR corresponds with a lower (or higher) release probability of OSNs in the olfactory system [22–25] as well as in other systems [26].

Fear conditioning via odor-foot shock pairing alters odor-induced responses imaged in the OB [4, 5, 7], suggesting a potential change in neurotransmitter release from OSNs. However, direct investigation into this question is hindered by technical challenges in unequivocal identification of conditioned presynaptic OSNs and their postsynaptic partners for electrophysiological recordings. Furthermore, a single odor typically activates multiple glomeruli and the activity of a single glomerulus may be influenced by other glomerular units via the OB network [27–29]. Mice in which OSNs with the M72 receptor coexpress the light-activated cation channel channelrhodopsin 2 (ChR2) can perceive optical stimulation of a single glomerulus [30]. These mice provide an ideal experimental model to address the question whether and how fear conditioning alters neurotransmitter release from genetically identifiable OSNs associated with a single glomerulus.

In this study, we conducted fear conditioning by optogenetically activating a single M72 glomerulus paired with foot shock and demonstrated that mice readily freeze to optical stimulation alone during fear retrieval. We then performed patch clamp recordings from genetically-labeled postsynaptic ET cells innervating M72 glomeruli and found a significant increase in OSN release probability in the conditioned glomerulus compared to control glomeruli (unstimulated or stimulated but without foot shock pairing). The increased release probability is accompanied by reduced GABA_B_ receptor expression in the conditioned glomerulus and is positively correlated with the animal’s freezing time. These results suggest that fear conditioning enhances peripheral olfactory inputs to the brain via GABA_B_-dependent modulation, which may directly contribute to the behavioral outcome.

## Results

### Pairing of monoglomerular activation and foot shock induces fear learning

To determine whether associative learning can be achieved by activating a single glomerulus, we used M72-ChR2/Vglut1-tdTomato mice (see Materials and Methods for details) in which olfactory sensory neurons (OSNs) expressing the M72 (or *Olfr* 160) receptor coexpress light-sensitive channelrhodopsin 2 (ChR2) and can be activated by blue light (473 nm) [30]. To optically stimulate the M72 glomeruli *in vivo*, an optical fiber with a diameter of 400 µm was implanted over a lateral M72 glomerulus of each mouse (Fig. 1A). We paired optical stimulation of a single glomerulus (conditioned stimulus) with mild foot shock (unconditioned stimulus) in a conditioning session and associative learning was assessed 24 hours later in a retrieval session (Fig. 1A, B). The conditioning session consisted of five trials of light-foot shock pairing with an inter trial interval of five minutes, and each pairing consisted of five 150 ms light stimulation at 0.5 Hz followed by 0.5 sec foot shock (0.5mA) (Fig. 1A). During the retrieval session, mice were placed in a novel environment. After three minutes (i.e., pre light-stimulation period), three trials of light stimulation (without food shock) were delivered with random intervals within 15 min while the animal was videotaped for behavioral assessment. The fear conditioned (NC) mice (n=12) showed robust freezing behavior, whereas the non-fear conditioned (NFC) mice (n=12) which underwent the same surgery and light stimulation procedure but without foot shock during the conditioning session did not (Fig. 1C). These results suggest that aversive learning can be achieved by optogenetic activation of a single glomerulus paired with foot shock.

**Figure 1.**
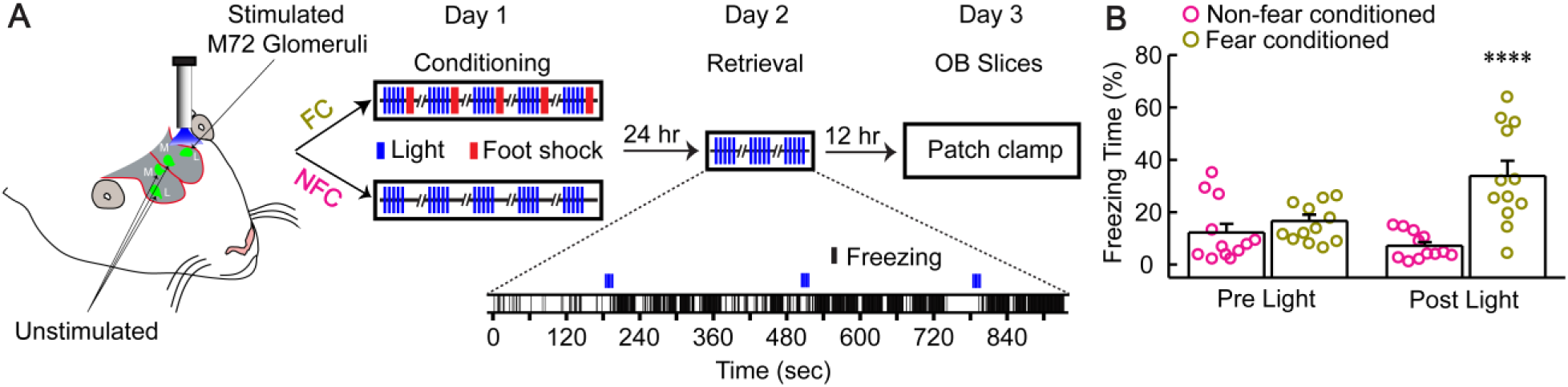
Aversive learning can be achieved by optogenetic activation of a single glomerulus paired with foot shock. **A**, Schematic drawing of a mouse head showing an optical fiber on one of the four M72 glomeruli and fear conditioning and retrieval paradigm in M72-ChR2/Vglut1-tdTomato mice. Blue vertical bars represent light stimulation and red bars foot shock. Inset, a representative retrieval session showing freezing bouts upon light stimulation previously paired with foot shock. **B**, During retrieval sessions, fear-conditioned but not control mice spent more time freezing post light-stimulation than pre light-stimulation (One Way ANOVA with Tukey correction, F_(3, 47)_=11.58, p=1.0E^−5^. FC pre light 15.6±2.1%, FC post light 33.8±5.4%, NFC pre light 12.2±3.3%, NFC post light 7.2±1.5%).

### Release probability of M72 OSNs is assessed in postsynaptic ET cells

In order to investigate how fear conditioning alters neurotransmitter release from OSNs, we used ET cells in the glomerular layer as the postsynaptic readout of transmitter release from M72 OSNs for several reasons. First, mitral cells are located farther away from the glomerulus and may receive weaker and indirect inputs from OSNs [31]. Second, in the glomerular layer, there are multiple types of inhibitory PG cells which receive either direct or indirect inputs from OSNs [14, 28]. Third, ET cells, a major excitatory element in the glomerular layer, receive strong monosynaptic inputs from OSNs [31, 32] and are readily identifiable via genetic labeling inM72-ChR2/Vglut1-tdTomato mice (Fig. 2A; See Materials and Methods for details).

**Figure 2.**
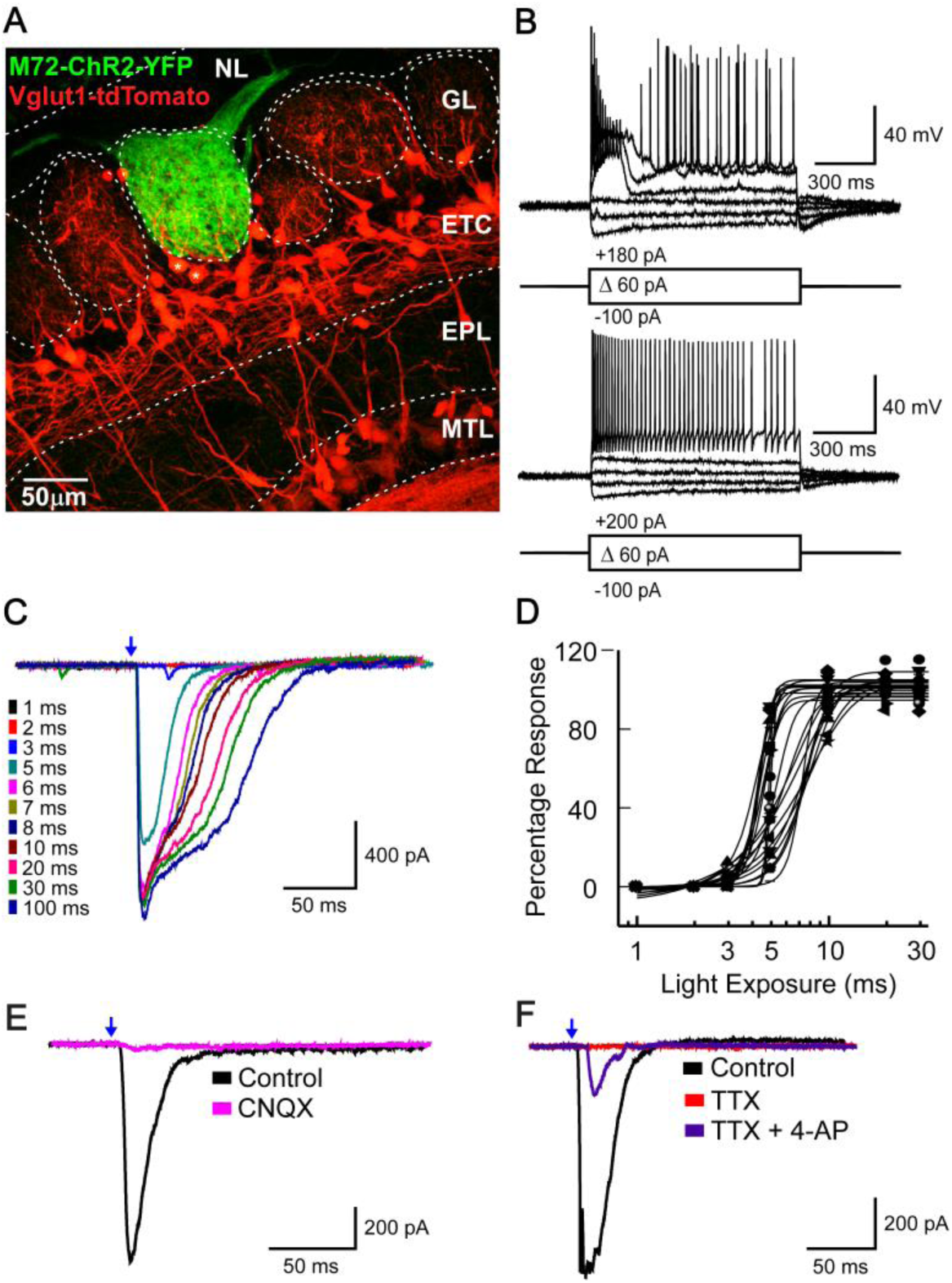
Transmitter release at M72-ChR2 glomeruli is assessed by measuring light-evoked EPSCs from Vglu1-tdTomato+ external tufted cells. **A**, Confocal image showing an M72-ChR2 glomerulus with Vglut1-tdTomato+ cells in different layers of the olfactory bulb. * ET neurons associated with the M72 glomerulus. **B**, Representative firing patterns demonstrating two subtypes of external tufted cells associated with M72 glomeruli under current clamp mode. **C**, Light-evoked EPSCs from an external tufted cell increase with stimulation duration. Note that the maximum peak current was achieved when the stimulation duration > 6 ms in this cell. **D**, The peak EPSCs were plotted against the stimulation duration from multiple cells and fitted with the Boltzmann function for sigmoid curves. For clarity, only a subset of the recorded cells (n=21) are included. **E,** Light-evoked EPSCs were blocked by bath perfusion of CNQX (25 μM). **F,** Light-evoked EPSCs were blocked by bath perfusion of TTX (0.5 μM) and reappeared after addition of 4-AP (1mM), suggesting monosynaptic sensory neuron inputs.

Approximately 12 hours after the retrieval session, the OB slices with “green” M72 glomeruli were selected for patch clamp recordings. In the glomerular layer, we specifically targeted Vglut1-tdTomato+ ET cells with discernible primary dendrites innervating the M72 glomeruli (Fig. 2A). Consistent with a previous study [33], two types of firing patterns were observed upon current injection: initial bursting (20 out of 64 or 31.3%; Fig. 2B, upper panel) and adapting (44 out of 64 or 68.8%; Fig. 2B, lower panel). Synaptic connection of a recorded ET cell with a M72-ChR2 glomerulus was confirmed by robust light-evoked excitatory postsynaptic currents (EPSCs) with little jitter (Fig. 2C). The light evoked EPSCs increased with the pulse length and reached the maximum peak current typically at ~10 ms (Fig. 2C, D). To compare OSN release properties across multiple glomeruli from different animals, we tested a series of pulse lengths to determine the minimum stimulus duration that evoked the maximum peak current in a given ET cell. This stimulus duration was then used for the release probability assessment in that cell. As expected, the light-evoked EPSCs were nearly eliminated by CNQX (25 µM; Control: 1473.3 ± 431.5 pA, CNQX: 22.7 ± 3.7 pA, and Washout: 770.7 ± 152.7 pA; n=4), an AMPA receptor antagonist, confirming glutamate mediated transmission (Fig. 2E). Light-evoked EPSCs were blocked by tetrodotoxin (TTX; 0.5 µM) and reappeared after application of TTX plus 1 mM 4-AP, a potassium channel blocker causing action potential independent release (n=3 out of 3), confirming monosynaptic connection (Fig. 2F).

### Fear conditioning increases release probability of OSNs

To assess transmitter release from OSNs, we measured the paired pulse ratio (PPR) [22–25] of light-evoked EPSCs in the ET cells innervating the M72 glomeruli with varying intervals (100, 200, and 300 ms). Compared to unstimulated M72 glomeruli in FC mice, the PPR of ET cells innervating the light-stimulated glomeruli was significantly reduced (Fig. 3A, B, E). Because the peak amplitude of the first EPSC (i.e., evoked by the first light pulse) was not significantly different among these two groups (FC-stimulated: 1152.94±161.93 pA; FC-unstimulated: 1137.29±117.75 pA; student t test, p=0.939), the reduced PPR was mainly due to the reduced amplitude of the second EPSC, indicating an increased release probability. To rule out that the increased release probability was merely due to light activation rather than fear conditioning, we performed the same comparison in NFC mice with light stimulation but without foot shock. In these NFC control mice, the PPR of ET cells was similar between light-stimulated and unstimulated glomeruli (Fig. 3C-E).

**Figure 3.**
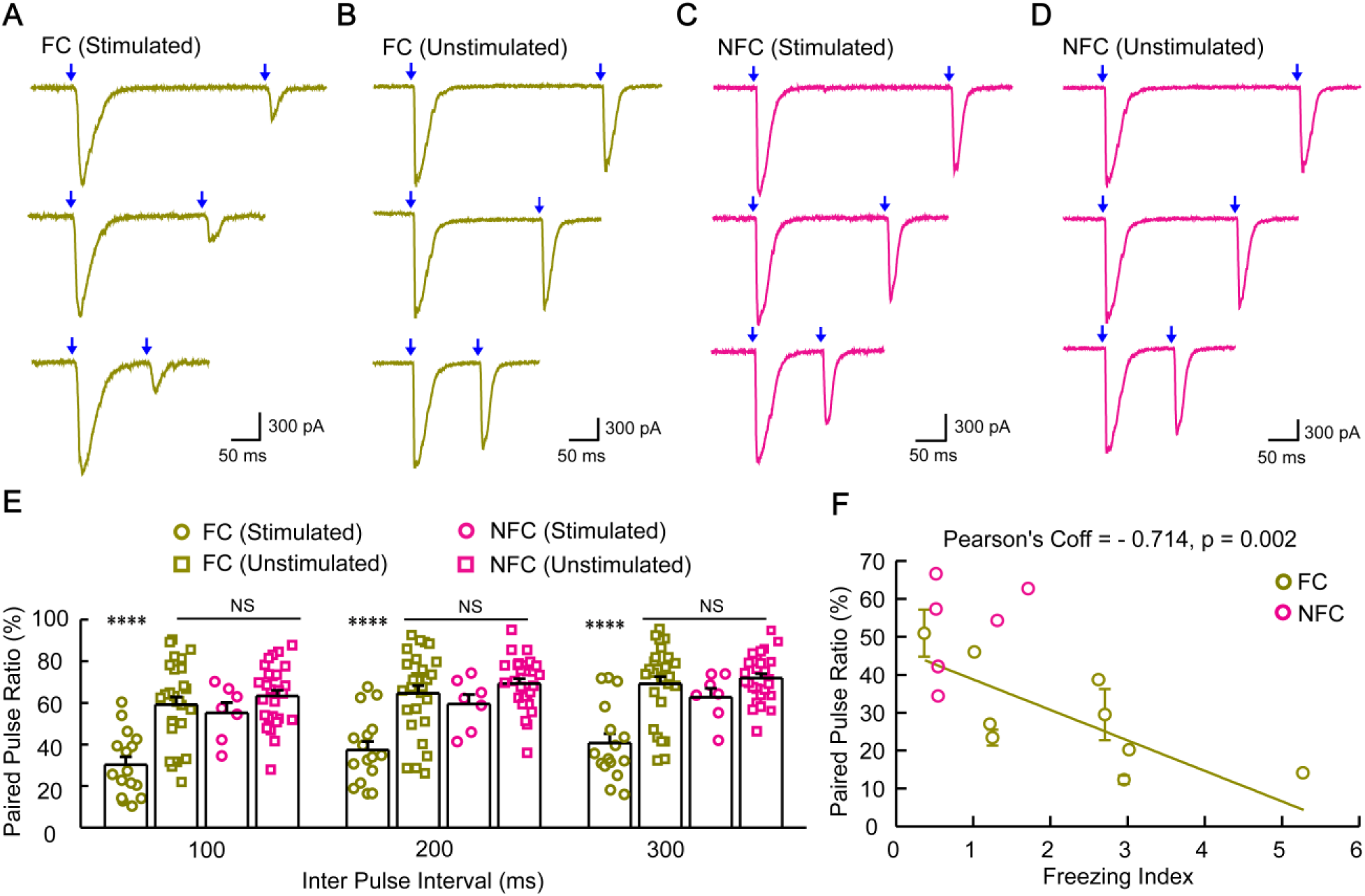
Fear conditioning increases the release probability of M72 OSNs. **A-D**, Light-evoked EPSCs from ET cells innervating different M72 glomeruli: stimulated (A, C) and unstimulated (B, D) in fear conditioned (FC) and non-fear conditioned (NFC) mice, respectively. Holding potential = −70 mV under voltage clamp mode. Blue arrows indicate onset of optical stimulation of associated M72 glomeruli and the inter-pulse interval (100, 200, or 300 ms) is defined from the end of the first pulse to the start of the second pulse. **E**, Summary of the PPR at varying intervals from ET cells innervating the four types of glomeruli: FC stimulated (100 ms interval: 30.3±3.8; 200 ms: 37.4±4.1; 300 ms: 40.7±4.5), FC unstimulated (100 ms: 59.2±3.7; 200 ms: 64.7±3.7; 300 ms: 69.2±3.4), NFC stimulated (100 ms: 55.2±4.9; 200 ms: 59.4±4.6; 300 ms: 62.7±4.3 and NFC unstimulated (100 ms: 63.3±2.8; 200 ms: 69.2±2.4; 300 ms: 71.9+2.2). One Way ANOVA with Tukey correction:100 ms: F_(3, 77)_=14.75, p=1.3E^−7^; 200 ms: F_(3, 77)_=14.24, p=2.0E^−7^; 300 ms F_(3, 77)_=15.53, p=6.2E^−8^. Each point represents the mean value of four to six trials from a single ET cell. Results from ET cells with different firing patterns were similar and thus grouped together. **F**, Relationship between the PPR (inter-pulse interval of 100 ms) and freezing behavior. In some mice, multiple ET cells associated with one glomerulus were recorded and the PPR was reported as mean ± SEM (with error bars) for that mouse.

As FC mice showed individual differences in the freezing time as well as in the release probability from the conditioned M72 glomeruli, we asked whether these two parameters were correlated. We quantified freezing behavior by calculating a freezing index, defined as percent freezing time post light-stimulation/percent freezing time pre light-stimulation during the retrieval session for each mouse. Out of the nine FC mice in which the PPR was successfully measured from the ET cell(s) associated with the light-stimulated M72 glomeruli, the PPR was negatively correlated with the freezing index (Fig. 3F), suggesting a positive correlation between the release probability and freezing time. In comparison, the five NFC mice showed little freezing and ET cells associated with the light-stimulated M72 glomerulus had high PPRs (Fig. 3F). These results suggest that fear conditioning increases the release probability from OSNs, which may contribute to the behavioral output.

Since we measured OSN transmitter release from the postsynaptic ET cells, the observed PPR changes could be due to pre- or postsynaptic mechanisms. To differentiate these two possibilities, we monitored the frequency and amplitude of spontaneous EPSCs (sEPSCs) in the ET cells innervating the M72 glomeruli from FC and NFC mice. While the amplitude of sEPSCs was similar among the four types of glomeruli (FC stimulated, FC unstimulated, NFC stimulated and NFC unstimulated), the frequency was significantly higher in the FC stimulated glomeruli (Fig. 4), supporting a presynaptic change in the OSN release properties.

**Figure 4.**
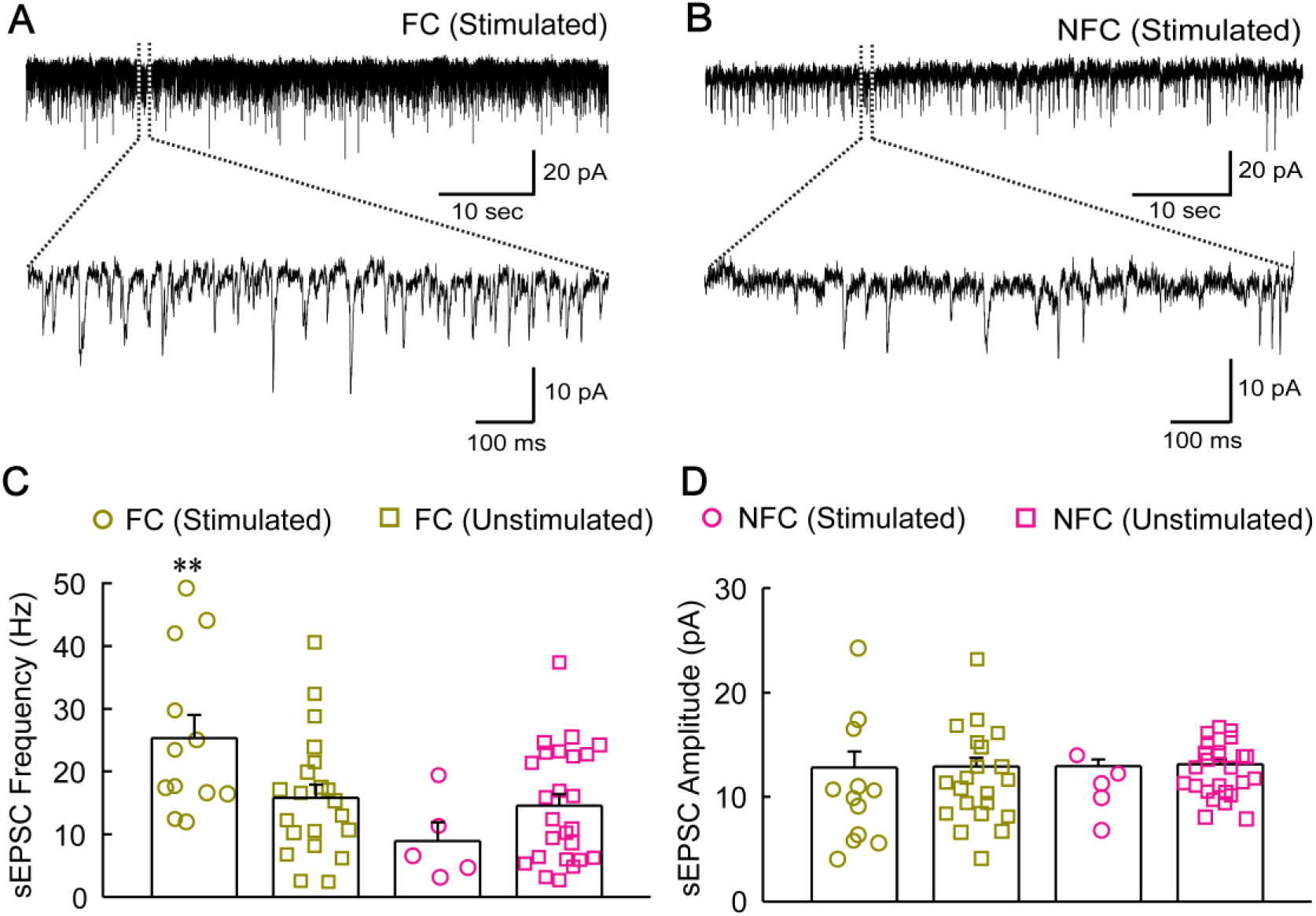
Fear conditioning increases the frequency but not amplitude of sEPSCs in the postsynaptic ET cells. **A-B**, Representative sEPSC traces recorded from an ET cell associated with the stimulated glomerulus in an FC (A) or NFC (B) mouse. Holding potential = −70 mV under voltage clamp mode. **C**, The frequency of sEPSCs from ET cells associated with different glomeruli: FC stimulated (25.3±3.7 Hz, n=12), FC unstimulated (15.8±2.1 Hz, n=21), NFC stimulated (8.9±3.0 Hz, n=5) and NFC unstimulated (14.6±1.8 Hz, n=25). There is an increase in the ET cells associated with the FC stimulated M72 glomeruli compared to the other three groups: F_(3, 62)_=4.47; p=0.006 (One Way ANOVA with Tukey correction). **D**, The amplitude of sEPSCs from the ET cells associated with different glomeruli: FC stimulated (12.8±1.5 pA, n=12), FC unstimulated (12.9±0.8 pA, n=21), NFC stimulated (12.9±0.7 pA, n=5), and NFC unstimulated (13.1±0.5 pA, n=25). There is no significant difference: F_(3, 62)_= 0.02, *P*=0.99 (one-way ANOVA with Tukey correction).

### Presynaptic GABA_B_ receptors may be involved in altered OSN release probability

A number of neurotransmitters and other molecules (GABA, dopamine, cyclic nucleotides, Zinc, serotonin, etc) in the glomerular layer have been implicated in modulating transmitter release from OSNs [<u>reviewed in 18</u>]. Among these possible mechanisms, the presynaptic GABA_B_receptor stands out as the most promising candidate as it has been shown to temper glutamate release from OSNs in a glomerulus-specific manner [12–17]. We hypothesized that GABA_B_ receptors are involved in fear learning induced transmitter release changes from OSNs associated with the conditioned glomerulus. Because GABA_B_ receptors are predominantly expressed in OSN axon terminals within glomeruli [34, 35], we examined the expression pattern of the GABA_B_ receptor 1 (R1) via antibody staining in the glomerular layer of FC and NFC mice. In the FC mice, the fluorescent intensity level (see Materials and Methods) of the stimulated M72 glomeruli was significantly lower than the unstimulated glomeruli in the same images (Fig. 5A-C). This difference was not observed in the NFC mice (Fig. 5C). These findings suggest that downregulation of GABA_B_ receptor expression may play a role in the increased release probability of OSNs after fear conditioning.

**Figure 5.**
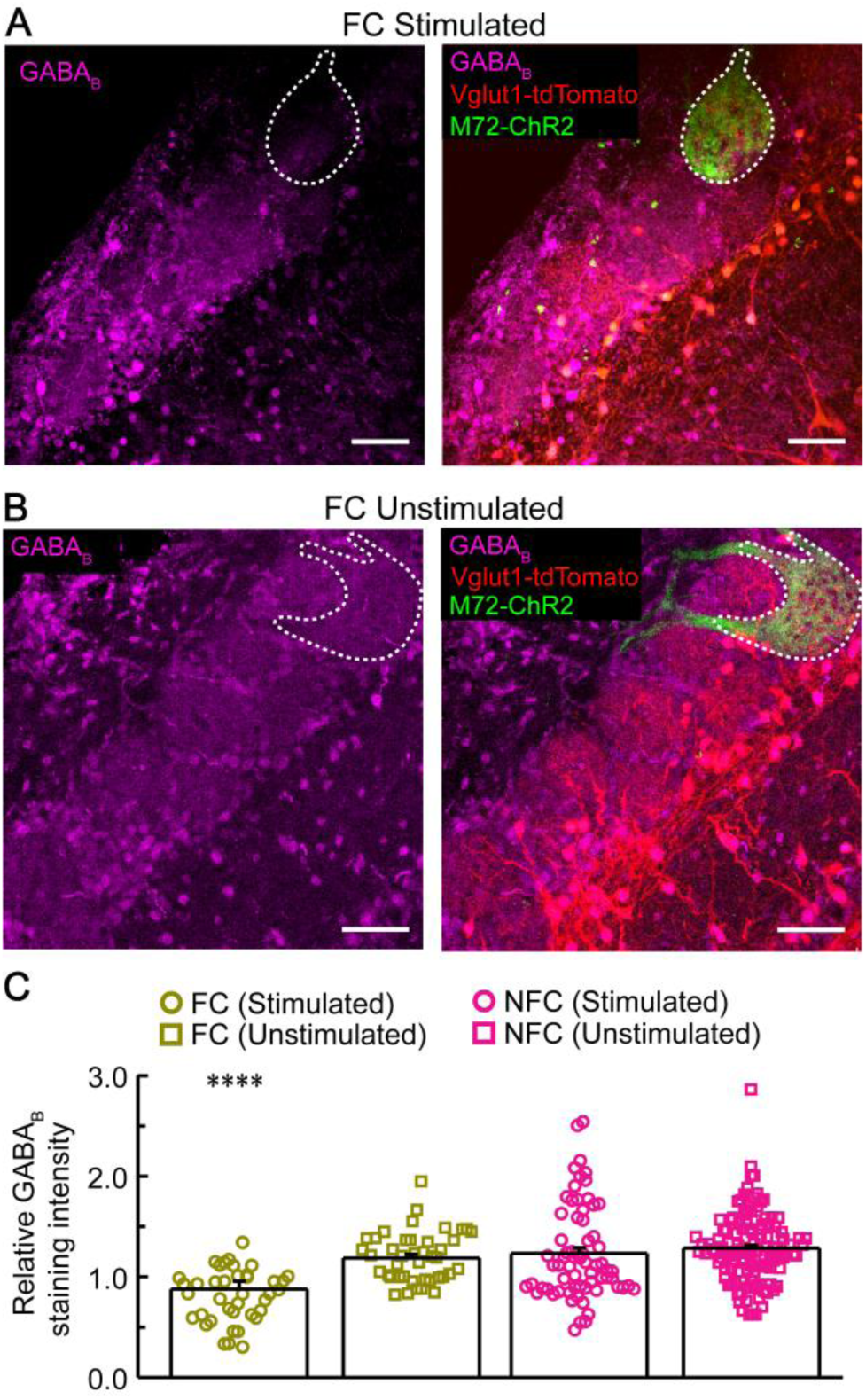
Fear conditioning decreases GABA_B_ receptor expression in the stimulated glomeruli. **A-B**, Confocal images of OB slices immuostained with GABA_B_ R1 antibody on the FC stimulated (A) and unstimulated side (B). Scale bars = 50 µm. M72 glomeruli are outlined by the dotted contour. **C**, Relative fluorescent levels of GABA_B_ antibody staining of M72 glomeruli in FC and NFC mice. Each dot represents the relative fluorescent intensity level of a neighboring glomerulus compared to the M72 glomerulus in the same image. We used the intensity ratio of M72/non-M72 glomerulus so that a lower expression level in the M72 glomerulus would be reflected as a smaller value. Fluorescent intensity of stimulated M72 glomeruli of FC mice was significantly lower than the control M72 glomeruli (the unstimulated glomeruli of FC mice, and both stimulated and unstimulated glomeruli of NFC mice) as compared to their neighboring glomeruli (FC mice: stimulated side 0.87±0.08, n=40 glomeruli vs FC-unstimulated side: 1.18±0.03, n=42 from four mice; NFC mice: stimulated side: 1.22±0.05, n=67 vs NFC-unstimulated side: 1.27±0.03, n=117 from four mice). One Way ANOVA with Tukey correction: F_(3, 265)_=10.31, p=1.9E^−6^.

If GABA_B_ receptors are indeed involved, activation or inactivation of these receptors would alter transmitter release from OSNs, which has been supported by previous studies using odor or olfactory nerve stimulation combined with pharmacological manipulations [12–17]. Here we examined the effects of GABA_B_ receptor activation or inactivation on spontaneous and optogenetically-evoked synaptic transmission from the ET cells associated with the M72 glomeruli. In the OB slices under control conditions, sEPSCs were evident from these ET cells (Fig. 6A). Application of Baclofen (20 μM), a GABA_B_ agonist, significantly decreased the frequency of sEPSCs (Fig. 6A, B), while CGP 55485 (CGP for short; 10 μM), a GABA_B_ antagonist, had the opposite effects (Fig. 6A, B). In contrast, neither drug changed the sEPSC amplitude significantly (Fig. 6C), supporting a role of GABA_B_ receptors in presynaptic inhibition of OSN transmitter release. Consistent with this notion, Baclofen decreased while CGP increased the light-evoked EPSCs in the ET cells (insets of Fig. 6D). Baclofen decreased the peak amplitude of the response to the first light stimulation by 50.1 ± 5.7% (paired t test, p=1.2E^−8^). When applied alone, CGP increased the peak amplitude by 18.3±4.4% (paired t test, p=0.002).

**Figure 6.**
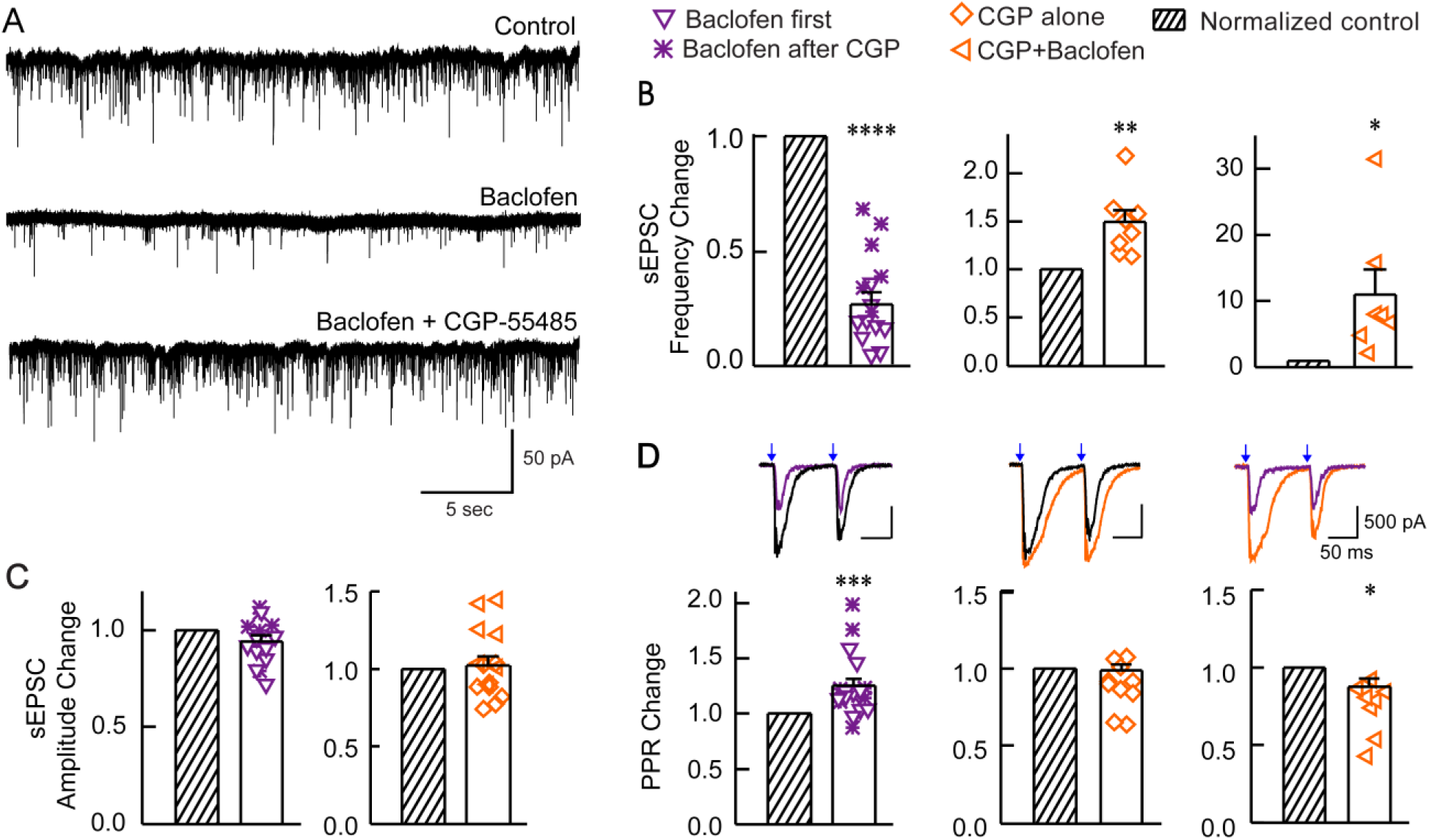
Pharmacological manipulation of GABA_B_ receptor activity changes the sEPSC frequency and OSN release probability measured in the ET cells. **A**, sEPSCs recorded from an ET cell under control conditions (upper panel), in the presence of Baclofen (20 µM; middle panel), and in the presence of Baclofen + CGP 55845 (10 µM; lower panel). Holding potential =-70 mV under voltage clamp mode. **B**, Summary data showing decreased or increased sEPSC frequency after Baclofen (left panel: paired t test, p=1.0E^−9^; n=15) or CGP application, respectively (middle panel for CGP alone: paired t test, p=0.004, n=8; and right panel for CGP + Baclofen: paired t test, p=0.039, n=7). In the left panel, two different symbols denote that Baclofen was applied first or after CGP, and the latter group showed less reduction, which may reflect incomplete washout of CGP. Each data point was analyzed from 2 min recording (3 min after drug application) and normalized to the value right before the drug application. **C**, There was no significant difference in the sEPSC amplitude after Baclofen or CGP (paired t test, Baclofen: p=0.081; CGP: p=0.670). **D**, Insets, Baclofen decreased while CGP increased light-evoked EPSCs in the ET cells. Baclofen increased the PPR of light evoked EPSCs in the ET cells associated with M72 glomeruli (left panel: paired t test, p=5.2E^−4^, n=20), which was reversed by CGP application (right panel: paired t test, p=0.032, n=8). No significant change in PPR was observed when CGP was applied alone (middle panel: paired t test, p=0.737, n=10).

When applied after Baclofen, CGP reversed the reduction caused by Baclofen. Furthermore, Baclofen increased the PPR of light-evoked EPSCs in the ET cells associated with the M72 glomeruli (Fig. 6D) supporting that activation of GABA_B_ receptors decreases the release probability of OSNs. The effect of Baclofen on the PPR was partially reversed by CGP, but application of CGP alone did not decrease the PPR as expected (Fig. 6D). One potential reason for this outcome is that we titrated the light stimulation duration to induce the maximum peak response in control conditions to measure PPRs, which leaves little room for the peak current to increase in the presence of CGP. In fact, CGP caused only modest increase in peak currents, but substantial increase in the area under light-evoked EPSCs reflecting the net charge carried by synaptic currents (inset of Fig. 6D). If we used the ratio of the area (second response/first response) to calculate PPR(area), CGP significantly decreased it (control 53.4± 3.6% vs CGP 43.1±3.0%; paired t test p=0.020). Taken together, these immunostaining and pharmacological experiments support a role of GABA_B_ receptors in modulating OSN transmitter release.

## Discussion

Here we demonstrate that stimulating a single glomerulus in the OB paired with foot shock is sufficient to induce fear learning in mice. This fear conditioning paradigm leads to enhanced neurotransmitter release from OSNs that project to the conditioned glomerulus, which is accompanied by reduced GABA_B_ receptor expression. Furthermore, the release probability of OSNs is positively correlated with the animal’s freezing time. This study provides the first direct evidence that fear conditioning enhances release probability from primary sensory neurons in any mammalian sensory system.

Many species including rodents rely mostly on olfactory cues to avoid danger and their olfactory systems display robust associative plasticity in various fear conditioning paradigms. Aversive learning induces plasticity not only in cortical and limbic areas [36–38], but also in early stages of the olfactory system [4–7, 39], suggesting that the same sensory stimulus sends different inputs to the brain depending on its predictive value. OSNs regenerate throughout life [40], and fear conditioning using an M71 ligand increases the number of M71 OSNs in the olfactory epithelium three weeks later [6]. This mechanism is unlikely to take place during short-term odor-foot shock pairing, which nevertheless increases conditioned odor-induced activity in the OB [4, 5, 7]. In genetically modified mice in which OSNs express the fluorescent exocytosis indicator synaptopHluorin [41], discriminative fear conditioning (CS+ with US and CS-without US) increases the glomerular responses to CS^+^ odors but not to CS-odors [5]. Similarly, classical fear conditioning (CS-US pairing) also increases the activity from mitral and tufted cells imaged at their dendrites in individual glomeruli [4, 7] as well as from periglomerular cells in the OB [39]. In this study, we provide direct electrophysiological evidence to support that fear conditioning changes peripheral olfactory inputs from OSNs to the OB neurons. Pairing optogenetic activation of a single glomerulus with foot shock leads to successful fear learning in M72-ChR2/Vglut1-tdTomato mice (Fig. 1). This is consistent with a previous study which shows that mice can perceive light activation of a single M72 glomerulus and detect small changes in odor-evoked glomerular spatial patterns [30]. This approach assigns learning to a single glomerulus with little interglomerular influence and permits further investigation of synaptic physiology in a cell-type specific manner, which would be otherwise impossible in classical odor-induced fear conditioning paradigms. The fear-conditioned M72 glomerulus and the associated postsynaptic ET cells are identifiable in OB slices, and the three unstimulated M72 glomeruli in the same animal or the stimulated M72 glomerulus from NFC animals serve as controls (Figs. 1, 2). Although our monoglomerular fear conditioning paradigm may not fully replicate the involvement of the OB circuit in olfactory learning, it offers unprecedented precision in revealing synaptic transmission changes from the conditioned OSNs. It is very likely that what we have learned here will also apply to classical olfactory learning because both approaches elicit robust freezing responses to cued stimuli and the effects of pharmacological manipulations are similar between olfactory nerve stimulation- and light-evoked EPSCs (see below).

Our patch clamp analysis reveals that the PPR of light-evoked EPSCs in the ET cells from the conditioned M72 glomeruli is significantly lower than the controls (unstimulated or stimulated without foot shock pairing) and the sEPSC frequency is higher (Figs. 3, 4), suggesting an increased release probability from presynaptic OSNs. This represents the first direct evidence that fear conditioning enhances release probability from primary sensory neurons in any mammalian sensory system. The yield of electrophysiological recordings from stimulated M72 glomeruli is relatively low as there is only one glomerulus/mouse with a small number of associated ET cells. Nonetheless, we were able to measure the PPR of light-evoked EPSCs in ET cells from nine FC mice and found a positive correlation between the release probability of OSNs and the freezing duration (Fig. 3). Further experiments are needed to determine to what extent the release probability change at the first synapse in the olfactory pathway contributes to the subsequent plasticity along the pathway and to the behavioral response in fear retrieval.

To shed light on the cellular mechanism underlying this fast form of neuroplasticity, we explored possible involvement of GABA_B_ receptors because presynaptic GABA_B_ receptors are known to temper transmitter release by suppressing calcium influx, activating inwardly rectifying K^+^ channels, and/or increasing the energy barrier for vesicle fusion [42, 43]. Here we show that the GABA_B_ receptor expression level in the conditioned M72 glomeruli is significantly lower than control M72 and other glomeruli (Fig. 5), consistent with the enhanced release probability and sEPSC frequency in the ET cells associated with the conditioned glomeruli (Figs. 3, 4). Fear conditioning may lead to internalization of GABA_B_ receptors in the presynaptic terminals of OSNs, as reported in other synapses [44]. It remains to be determined what signals cause the downregulation of GABA_B_ receptors in the conditioned glomeruli, but most likely it involves centrifugal inputs into the OB. For instance, the glomerular layer receives extensive top-down inputs from olfactory cortices and neuromodulatory centers (e.g., serotonin from raphe nuclei, acetylcholine from basal forebrain, and norepinephrine from locus ceruleus) [45–48].

We notice that the immediately adjacent glomeruli of the conditioned M72 glomerulus also show lower GABA_B_ receptor expression in contrast to the more uniform expression among the unstimulated and their neighboring glomeruli (Fig. 5). This finding suggests that the mechanisms that are involved in downregulating GABA_B_ receptors may not selectively target the conditioned glomerulus only, but rather a small area covering multiple glomeruli. Together with other fear conditioning induced changes in the OB circuit and neuromodulation [3, 5], this phenomenon may contribute to olfactory fear generalization [4, 39]. To test this possibility, future experiments would require activating adjacent glomeruli individually while recording from their postsynaptic cells.

Although the immunostaining experiments shown in Fig. 5 do not differentiate GABA_B_ receptors on the pre- or post-synaptic cells, previous studies using immuno-electron microscopy demonstrates that GABA_B_ positive neuropils within glomeruli are predominantly axon terminals of the olfactory nerve [34, 35]. These GABA_B_ receptors mediate presynaptic inhibition and suppress glutamate release from OSNs, supported by ex vivo as well as in vivo pharmacological studies [12–17]. In OB slices, the GABA_B_ receptor agonist Balcofen decreases olfactory nerve stimulation evoked EPSCs in both PG and ET cells, while the GABA_B_ antagonist CGP has the opposite effect [14]. Here we show that these pharmacological manipulations have similar effects on the evoked EPSCs in the ET cells upon optogenetic activation of OSNs associated with a single glomerulus (Fig. 6). In addition, Baclofen decreases and CGP increases the sEPSC frequency in the ET cells, supporting a presynaptic action of GABA_B_ receptors (Fig. 6).

The effects of these drugs on the PPR of the evoked EPSCs are more complicated. Activating GABA_B_ receptors by Baclofen increases the PPR mainly by reducing the response to the first stimulus, which is at least partially reversed by CGP (Fig. 6). The potential role of GABA_B_ receptors in mediating the OSN release probability is also supported by previous reports that CGP decreases the PPR evoked by olfactory nerve stimulation and measured in postsynaptic cells [14, 17]. However, in our slice recordings, blocking GABA_B_ receptors by CGP alone does not change the PPR. This could be due to the use of light stimulation that induces near maximum response leaving little room for the peak current to increase. When we calculate the PPR based on the net charge carried by EPSCs, CGP decreases the PPR(area). Additionally, this could be due to a relatively low activity of GABA_B_ receptors in the absence of agonists in control conditions. Moreover, our finding could mean that fear conditioning induced changes in light-evoked EPSCs involves both GABA_B_-dependent and GABA_B_-independent mechanisms, such as OSN synaptogenesis [49].

Olfactory aversive learning induced neuroplasticity has been reported in other species including humans. After aversive conditioning using two enantiomers indiscriminable to subjects, the evoked activity pattern in the piriform cortex becomes more distinct and the ability to discriminate the odors is enhanced [50]. It is plausible that plasticity observed in the piriform cortex reflects cumulative changes from the previous stages including the OSNs and the OB network as well as the cortical circuits.

What we have learned from the olfactory pathway may also apply to other sensory modalities as well. Presynaptic GABA_B_ receptors and local inhibitory neurons are known to modulate visual and auditory glutamatergic inputs at their first synapses in the brain [20, 21]. Fear learning based on visual or auditory cues may increase the release probability of the neurons that respond to the conditioned stimulus via downregulation of presynaptic GBAB_B_ receptor expression. Our study reveals fear conditioning induced neuroplasticity at the first synapse in the olfactory pathway, highlighting the ability of the brain to adjust sensory inputs based on predictive values.

### Material and Methods

#### Animals

M72-IRES-ChR2-YFP (in brief M72-ChR2) mice (JAX Stock No: 021206) in which OSNs expressing the M72 receptor coexpress fused ChR2-yellow fluorescent protein were provided by Drs. Thomas Bozza and Dmitry Rinberg [30]. In order to achieve fluorescent labeling of the OB projection neurons for electrophysiological recordings, M72-ChR2 mice were crossed with Vglut1-Cre mice (Vglut1-IRES2-Cre-D; JAX Stock No: 023527) [51] and the Cre-dependent tdTomato reporter mice (Rosa26-CAG-LSL-tdTomato-WPRE or Ai9 line; JAX Stock No: 007905) [52] to obtain triple transgenic M72-ChR2^+/+^;Vglut1^Cre/+^;Rosa^tdTomatof/+^ mice (in brief M72-ChR2/Vglut1-tdTomato) for experimental use. All mice were housed under 12 hr light-12 hr dark cycles with food and water ad libitum in a temperature- and humidity-controlled animal facility. Mice were randomly assigned for control or experimental groups. All behavioral and recording procedures were performed during the light phase of the cycle. Both male and female mice (2-3 months old, unless otherwise stated) were used. All procedures were approved by the University of Pennsylvania Institutional Care and Use Committee.

### Surgical Implantation

Mice were anesthetized via isoflurane at 3% (vol/vol) and secured in a stereotaxic system (Model 940, David Kopf Instruments). Isoflurane levels were maintained at 1.5-2% for the remainder of the surgery. Body temperature was maintained at 37 °C with a temperature control system (TC-1000, CWE Inc.). The overlying bone of the left olfactory bulb was thinned until the blood vessels over the OB were clearly visible and the lateral M72 glomerulus was located under a fluorescent dissecting microscope (Leica M80). A fiber optic cannula stub (400 µm core, 2.5 mm fiber length, Thor Labs) was positioned above the glomerulus and fixed in place with Kwik-Sil adhesive (World Precision Instruments) and dental cement. After surgery, mice were caged individually and were allowed to recover for 1 week prior to fear conditioning.

### Fear Conditioning and Retrieval

In the fear conditioning session (Day 1), mice were acclimatized to the foot shock chamber (Modular Chamber, ENV-307A, Med Associates Inc.) for at least one hour. Fear conditioning was performed by pairing optical stimulation (conditioned stimulus or CS: five 150 ms pulses at 0.5 Hz, with each light pulse mimicking the length of an inhalation) with mild foot shock (unconditioned stimulus or US: 0.5 sec at 0.5 mA). The onset of the US coincided with the offset of the CS and such pairing repeated five times with an inter-trial interval of five minutes (Fig. 1A). Optical stimulation was delivered from a 473nm laser (SLOC Lasers, BL473T8-150FC) through a fiber optic rotary joint patch cable (Thor Labs). The laser power was adjusted to 50 mW/mm^2^ measured from the optic fiber tip and the timing was controlled by an arbitrary waveform generator (Agilent 33201A). In the fear retrieval session 24 hours later (Day 2), mice were put into a novel chamber (7 1/2” x 11 1/2” x 5”). After 3 min of free exploration period, three trials of light stimulation (five 150 ms pulses at 0.5 Hz) at a random interval of 3 to 5 minutes were presented without foot shock within 15 min. To prevent mice from using the leaked light as a visual cue for associative learning, bright blue LED was positioned over the fear conditioning and retrieval chamber to mask the dull leaked light. Freezing behavior was scored manually by visual inspection of recorded videos and animals were considered to be freezing if no movement was detected for 1?sec by an experimenter blind to experimental conditions.

### Patch Clamp Recording

Approximately 12 hr after the retrieval session, the mice were sacrificed for electrophysiological recordings in acute brain slices. Mice were deeply anesthetized with ketamine-xylazine (200 and 15 mg/kg body weight, respectively) and decapitated. The brain was dissected out and immediately placed in ice-cold HEPES buffer. Using a Leica VT 1200S vibratome, relatively thin coronal OB slices (thickness=130 μm) were cut to facilitate visualization of the M72 glomeruli (diameter ~70 μm) and accessibility of surrounding cells. Slices were incubated in oxygenated artificial cerebrospinal fluid (ACSF) containing the following (in mM): 126 NaCl, 2.5 KCl, 2.4 CaCl_2_, 1.2 MgSO_4_, 11 D-glucose, 1.4 NaH_2_PO_4_ and 25 NaHCO_3_ (osmolality ~305 mOsm and pH 7.4, bubbled with 95% O_2_-5% CO_2_) for 1 hr at 31°C and kept in oxygenated ACSF in room temperature thereafter. Before recording, slices were transferred to a recording chamber and continuously perfused with oxygenated ACSF. M72-ChR2-YFP glomeruli and Vglut1-tdTomato^+^ cells were visualized with a 40× water-immersion objective under an Olympus BX51WI upright microscope equipped with epifluorescence. Blue light used to activate ChR2 in the OB slices was provided by a Lambda DG-4 (Sutter Instrument) through a band pass filter (450 to 490 nm).

Whole-cell patch-clamp recordings were made in both current and voltage-clamp mode. Recording pipettes were made from borosilicate glass (GC210F-10; Harvard Apparatus) with a Flaming-Brown P-97 puller (Sutter Instruments; tip resistance 5–8 M?). The pipette solution contained the following (in mM): 120 K-gluconate, 10 NaCl, 1 CaCl2, 10 EGTA, 10 HEPES, 5 Mg-ATP, 0.5 Na-GTP, and 10 phosphocreatine. Electrophysiological recordings were controlled by an EPC-10 amplifier combined with Pulse Software (HEKA Electronic) and analyzed using Igor Pro and mini-analysis (Synaptosoft, Inc.). The signals were filtered at 2.9 kHz and acquired at 10 kHz. Pharmacological reagents including tetrodotoxin (TTX) citrate, 6-cyano-7-nitroquinoxaline-2,3-dione (CNQX), 4-Aminopyridine (4-AP), Baclofen (Sigma-Aldrich) and CGP 55845 (Tocris Bioscience) were bath perfused.

### Immunohistochemistry

After electrophysiological recordings, brain slices were fixed in 4% paraformaldehyde (Sigma) for 15-20 min at 4 °C. Sections were washed in 1% Triton X-100 and 1% Tween 20 in PBS (3X 10 minutes), blocked for 60 min in 1% Triton X-100 and 1% Tween 20 in PBS with 3% bovine serum albumin, and then incubated at 4 °C with the mouse anti-GABA_B_-R1 (1:500, H00002550-M01, ABNOVA) primary antibody in the same solution overnight. Immunofluorescence was achieved by reaction with the secondary antibody goat anti-mouse, Alexa Fluor 633 (A21052, Molecular Probes, Invitrogen) at 1:1000 overnight at 4 °C. Tissues were washed in 1% Tween 20 in PBS (3X 20 min) and mounted with Fluoromount-G (Southern Biotech). Fluorescent images were taken under a SP5/Leica confocal microscope with LAS AF Lite software. The fluorescent intensity of individual glomeruli was quantified from unprocessed images using ImageJ. For each image, the same-size circle of ~600 µm^2^ was placed in the middle of each glomerulus void of stained cell bodies and the fluorescent intensity within the circle was measured. After subtracting the background intensity (measured using the same circle placed in an area void of GABA_B_ staining), the relative intensity for each non-M72 glomerulus compared to the M72 glomerulus on the same image was calculated.

### Statistics

All data were expressed as mean ± S.E.M. Student *t-*test was used for statistical comparison of two groups and one-way ANOVA followed by post hoc Tukey test for multiple groups (ORIGIN 8.0). A *p* value *<*0.05 was considered to be significant (* p<0.05, ** p<0.01, *** p<0.001, and **** p<0.0001).

## Acknowledgements

We thank Drs. Thomas Bozza and Dmitry Rinberg for sharing the M72-IRES-ChR2-EYFP mouse line and genotyping protocol. This work was supported by the National Institute of Health (NIDCD R01 DC006213 to M.M. and F31 DC017054 to M.S. and NIMH T32 MH017168 to M.S.).

